# Orb prevents autophagy in the *Drosophila* germline through translational repression of *Atg12* mRNA

**DOI:** 10.1101/007971

**Authors:** Isabelle Busseau, Stéphanie Pierson, Dany Séverac, Christelle Dantec, Martine Simonelig

**Author notes:** Corresponding authors: Isabelle Busseau; Martine Simonelig.

## Abstract

*Drosophila* Orb, the homologue of vertebrate CPEB is a key translational regulator involved in oocyte polarity and maturation through poly(A) tail elongation of specific mRNAs. *orb* has also an essential function during early oogenesis which has not been addressed at the molecular level. Here, we show that *orb* prevents cell death during early stages of oogenesis, thus allowing oogenesis to progress. It does so through the repression of autophagy, by directly repressing, together with the CCR4 deadenylase, the translation of *Autophagy-specific gene 12* (*Atg12*) mRNA. The uncontrolled autophagy observed in *orb* mutant ovaries is reduced when *Atg12* mRNA levels are decreased. These results reveal a role of Orb in translational repression and identify autophagy as an essential pathway regulated by Orb during early oogenesis. Importantly, they also establish translational regulation as a major mode of control of autophagy, a key process in cell homeostasis in response to environmental cues.

## Introduction

The regulation of developmental processes and cellular activities largely relies on translational control of mRNAs, and an important mechanism of this regulation involves changes in mRNA poly(A) tail lengths (Weill et al., 2012). Cytoplasmic polyadenylation element binding (CPEB) proteins act sequentially in poly(A) tail shortening and lengthening through the recruitment of deadenylases and poly(A) polymerases (Richter, 2007). They bind to UA-rich short sequences, referred to as cytoplasmic polyadenylation elements (CPEs), located within 3’UTRs of their target mRNAs (Pique et al., 2008; Richter, 2007). CPEB1 has been mostly studied for its implication in vertebrate oocyte maturation, where it is involved in translational repression through the recruitment of PARN deadenylase in immature oocytes, and in translational activation through an interaction with Gld2 poly(A) polymerase during maturation (Igea and Mendez, 2010; Kim and Richter, 2006). The role of CPEBs in translational regulation in somatic tissues has also been established in various contexts, including the control of cell proliferation, senescence, tumor development, synaptic plasticity and glucide metabolism (Alexandrov et al., 2012; Bava et al., 2013; Fernandez-Miranda and Mendez, 2012; Ortiz-Zapater et al., 2012; Udagawa et al., 2012).

Orb is the *Drosophila* homologue of vertebrate CPEB1 and, consistent with this, it is involved in cytoplasmic polyadenylation and translational activation during oogenesis, for oocyte polarity and maturation (Benoit et al., 2008; Castagnetti and Ephrussi, 2003; Chang et al., 1999; Juge et al., 2002). However, while Orb plays an established role in the initial steps of egg chamber formation, this function is poorly understood (Christerson and McKearin, 1994; Huynh and St Johnston, 2000; Lantz et al., 1994).

Here we show that cell death is the major defect in *orb* mutant early ovaries. Developmental programmed cell death occurs at three specific stages in the female germline: in the newly formed cysts (region 2 of the germarium, Figure 1A), during mid-oogenesis (stages 7-8), and during late oogenesis (stages 12-13) (McCall, 2004; Pritchett et al., 2009). Germ cell death during early and mid-oogenesis is strongly enhanced by starvation (Drummond-Barbosa and Spradling, 2001), and strikingly does not depend on the usual apoptotic activators such as *hid*, *grim*, *reaper* (Foley and Cooley, 1998; Peterson et al., 2007; Pritchett et al., 2009), but involves autophagy (Barth et al., 2011; Nezis et al., 2009). In *Drosophila*, autophagy contributes to developmental cell death in several developmental processes through caspase activation and DNA fragmentation (Denton et al., 2012). A molecular link has been established between autophagy and cell death during late oogenesis as nurse cell death by DNA fragmentation depends on autophagic degradation of the inhibitor of apoptosis (IAP/dBruce) (Nezis et al., 2010).

**Figure 1.**
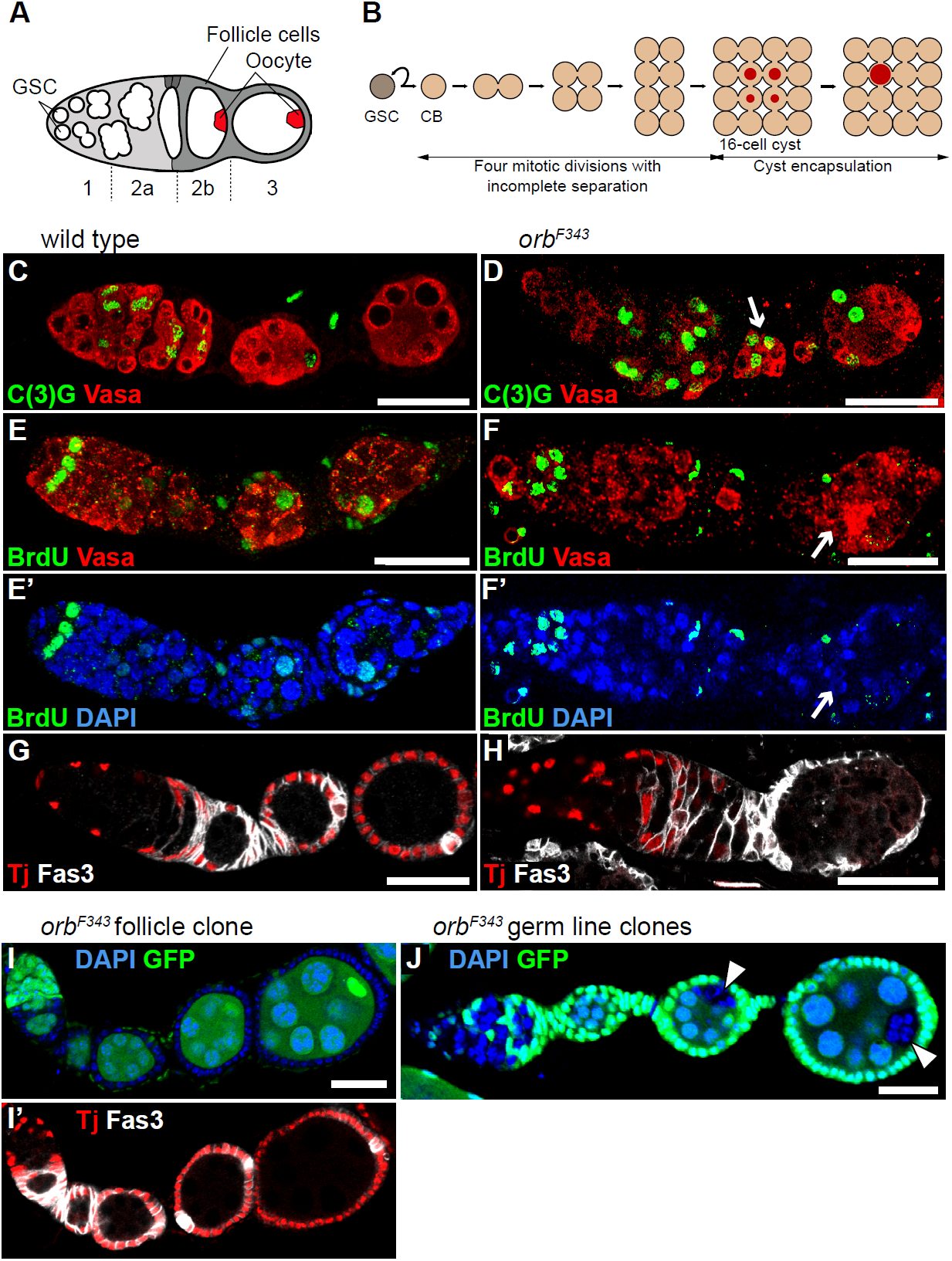
Cell cycle and meiosis defects in *orb* mutant germ cells. (A) Schematic drawing of a wild-type germarium. Germ cells are represented in white and the differentiated oocyte in red (GSC, germline stem cells). Somatic follicle cells are in grey. (B) Schematic drawing of the mitotic divisions producing a cystoblast (CB) from one GSC, and a 16 germline cell-cyst from one CB. Red dots indicate meiosis which initiates in four of the 16 germ cells within a cyst, and becomes restricted to the oocyte. (C-H) Defects in meiosis restriction and cell cycle in *orb*^*F343*^ germ cells. Wild type (left panels) and *orb*^*F343*^ (right panels) ovarioles stained with anti-Vasa (germ cell marker, red) and anti-C(3)G (green) (C, D); anti-Vasa (red) (E, F) or DAPI (E’, F’) and BrdU incorporation (green) (E-F’); anti-Tj (red) and anti-Fas3 (white) (G, H). White arrows indicate the lack of C(3)G restriction to one cell in (D) and the lack of BrdU incorporation in pseudo-cyst nuclei in (F, F’). (I, I’) *orb*^*F343*^ follicle clone labeled with DAPI (blue) and GFP (green) (I), and with anti-Tj (red) and anti-Fas3 (white) (I’). (J) *orb*^*F343*^ germline clones labeled with DAPI (blue) and GFP (green). White arrowheads indicate *orb*^*F343*^ germ cells. Scale bars: 20 μm.

Here, we show that *orb* prevents autophagy and cell death during early stages of oogenesis, and identify *Autophagy-specific gene 12* (*Atg12*) mRNA as a direct target of translational repression by Orb.

## Results and Discussion

### Early defects in *orb* mutant ovaries

We sequenced *orb^F343^* (Lantz et al., 1994) and *orb^36-53^* (Morris et al., 2003), two strong or null *orb* alleles and found that both have premature stop codons ( Figure S1A, B). For all subsequent analyses, we used *orb^F343^* which had the most upstream stop codon. We used meiosis and cell cycle markers to further address the early germ cell defects reported previously (Huynh and St Johnston, 2000; Lantz et al., 1994). Oogenesis stops as pseudo-cysts just after the germarium in *orb^F343^*. In these pseudo-cysts, expression of C(3)G, a component of the synaptonemal complex, was generally maintained in several cells, reflecting a defect in meiosis restriction and, hence, oocyte determination (Figure 1C, D) (Huynh and St Johnston, 2000). Incorporation of bromodeoxyuridine (BrdU) was used to monitor germ cell DNA replication. BrdU incorporation that is detected in region 1 of wild-type germarium was not affected in *orb* mutants (Figure 1E-F’). In contrast, the BrdU incorporation found in wild-type endoreplicating nurse cells was never detected in *orb* mutant pseudo-cysts (Figure 1E-F’), suggesting that the germ cells in these pseudo-cysts had not entered the nurse cell fate.

In the wild type, follicle cells encapsulate individual cysts to produce egg chambers. They express the transcription factor Traffic Jam (Tj) and Fasciclin 3 (Fas3) and, as they mature, Fas3 is down-regulated except in polar cells (Figure 1G). In *orb^F343^* mutant ovaries, the follicle cells did not express Tj and failed to down-regulate Fas3, suggesting a defect in follicle cell maturation (Figure 1H). We used FLP-mediated FRT recombination to generate *orb* mutant cell clones and investigate a potential intrinsic function of *orb* in follicle cells. Ovaries with *orb* mutant follicle cells produced normal egg chambers (Figure 1I, I’), demonstrating that *orb* function was not required in the follicle cell lineage. Analysis of *orb* mutant germline clones showed that *orb* was dispensable in the germline stem cells for their self-renewal, division rate and differentiation (Figure S1C-F), consistent with *orb* function being downstream of these events. *orb* mutant germ cells were co-encapsulated with wild-type cysts in compound egg chambers (70%, n = 237) (Figure 1J).

We conclude that *orb* is required in the germ cells for meiosis restriction and endoreplication of nurse cell nuclei. Defects in these processes prevent germ cell differentiation into oocyte and nurse cells and affect follicle cell maturation non-autonomously.

### Germ cell death is a major defect in *orb* mutant ovaries

DAPI staining of *orb* mutant germline clones revealed pycnotic nuclei that can indicate cell death. We therefore used anti-cleaved caspase 3 and TUNEL assays to record potential cell death in *orb* mutant ovaries. The staining with both markers revealed cell death in *orb* mutant germ cells (89% (n = 140) and 69% (n = 81) of pseudo-cysts marked with anti-cleaved caspase 3 and TUNEL, respectively) (Figure 2A-D). We used the *UAS/Gal4* system to overexpress the known caspase inhibitor DIAP1 in the germline (Mazzalupo and Cooley, 2006; Peterson et al., 2003). DIAP1 expression reduced the levels of cleaved caspase 3 and TUNEL staining in *orb* mutant germ cells (Figure 2E-F), and strikingly, rescued the formation of egg chambers in 23 to 27% of ovarioles, with follicle cells expressing Tj and Fas3 normally (Figure 2F', G). This showed that germ cell death contributed to the early oogenesis arrest in *orb* mutants. Oogenesis did not progress further in these rescued egg chambers, consistent with Orb regulating other processes in the ovary.

**Figure 2.**
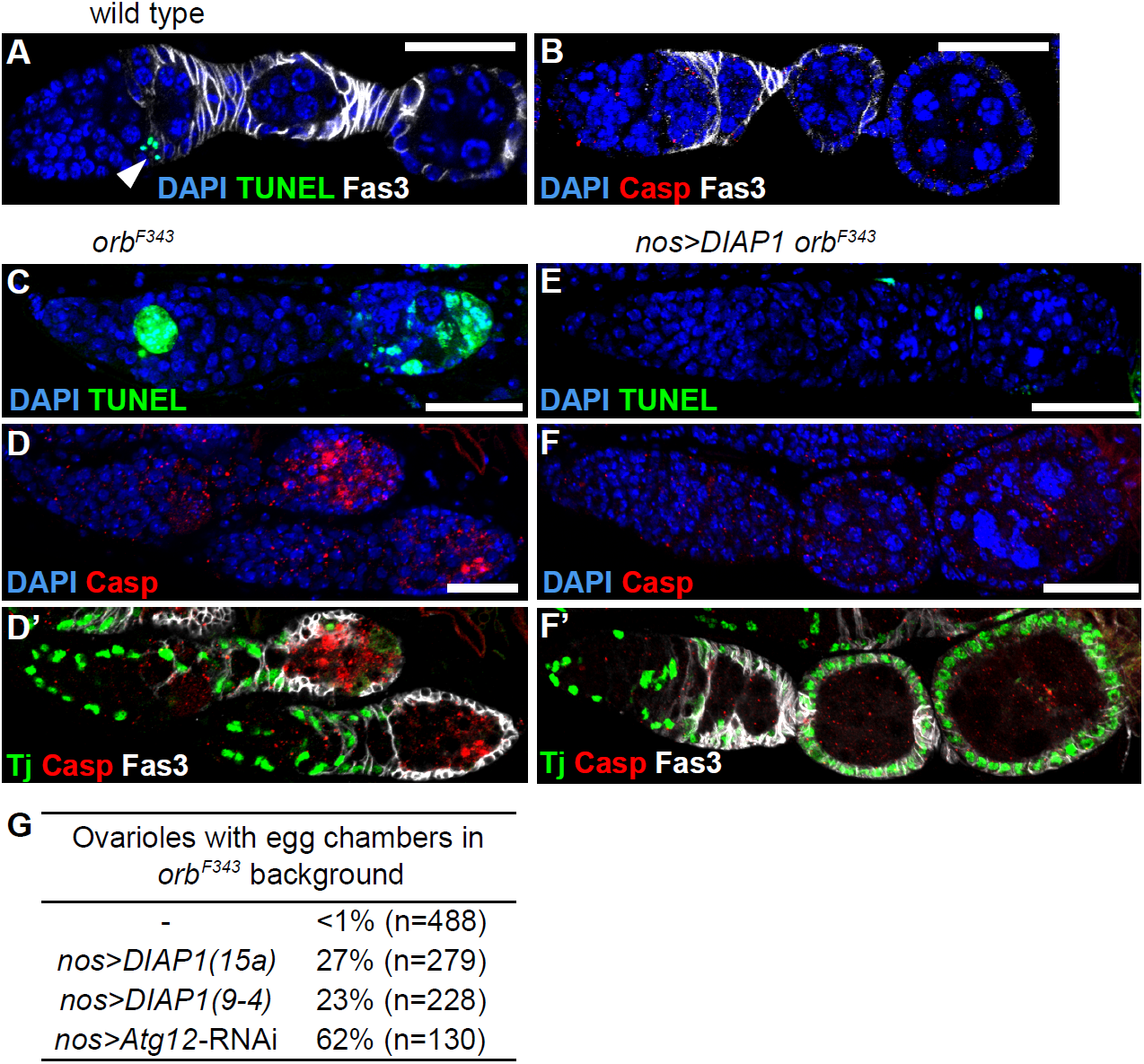
Cell death in *orb* mutant ovaries. (A, B) Wild-type ovarioles labeled with DAPI (blue), TUNEL (green), anti-cleaved caspase 3 (Casp, red) and anti-Fas3 (white). The white arrowhead indicates TUNEL staining in follicle cells. (C-F’) *orb*^*F343*^ (C-D’) and *UASp-DIAP1/+; nos-Gal4 orb*^*F343*^/*orb*^*F343*^ (*nos > DIAP1 orb*^*F343*^) (E-F’) ovarioles labeled with DAPI (blue) and TUNEL (green) (C, E); DAPI (blue), anti-cleaved caspase 3 (Casp, red) (D, F); and anti-Tj (green), anti-cleaved caspase 3 (Casp, red) and anti-Fas3 (white) (D’, F’). Scale bars: 20 μm. (G) Quantification of ovarioles containing egg chambers in *orb*^*F343*^ either expressing DIAP1 (*UASp-DIAP1/+; nos-Gal4 orb*^*F343*^/*orb*^*F343*^ (*nos > DIAP1*)) or *Atg12-*RNAi (*UASp-TRIP-Atg12 orb*^*F343*^/*nos-Gal4 orb*^*F343*^ (*nos > Atg12-RNAi*)).

### Identification of mRNA targets of Orb

To address whether *orb* could regulate cell death directly we performed Orb RNP immunoprecipitation-microarray (RIP-Chip) analysis (Keene, 2007) to identify mRNAs associated with Orb. We used either mature ovaries or early ovarian stages dissected from newly eclosed females (germarium to stage 8) (Figure S2). Significance analysis of microarrays (SAM) with a false discovery rate (FDR) of 0.01 then identified 421 and 603 mRNAs that were enriched at least 1.5 fold in Orb RIP from early and mature ovaries, respectively, compared to mock RIP (Figure 3A, Table S1). Gene ontology (GO) term enrichment analysis using DAVID with a p-value <0.05 (Benjamini corrected) identified the terms “translation”, “cell cycle” and “mitochondria” as enriched among the mRNAs present in Orb RIP (Figure 3B). While CPEs have been defined in *Xenopus* (Pique et al., 2008), they remain uncharacterized in *Drosophila*. We used the software designed to identify CPEs in *Xenopus* (Pique et al., 2008) to identify those in mRNAs from Orb RIP. CPEs were not found enriched in Orb RIP mRNAs compared to mRNAs expressed in ovaries (16%, versus 22% in 6614 mRNAs expressed in ovaries from FlyBase). This could indicate that the software did not reveal all Orb binding motifs. It appears unlikely because Orb possesses most aminoacids shown in CPEB1 to be involved in the interaction with the CPE (Afroz et al., 2014). It is important to note that Orb is part of large ribo-nucleoprotein complexes, where it is associated with many other RNA binding proteins (Weill et al., 2012). It is therefore very likely that a proportion of mRNAs in Orb RIP could coprecipitate through interactions with these other RNA-binding proteins.

**Figure 3.**
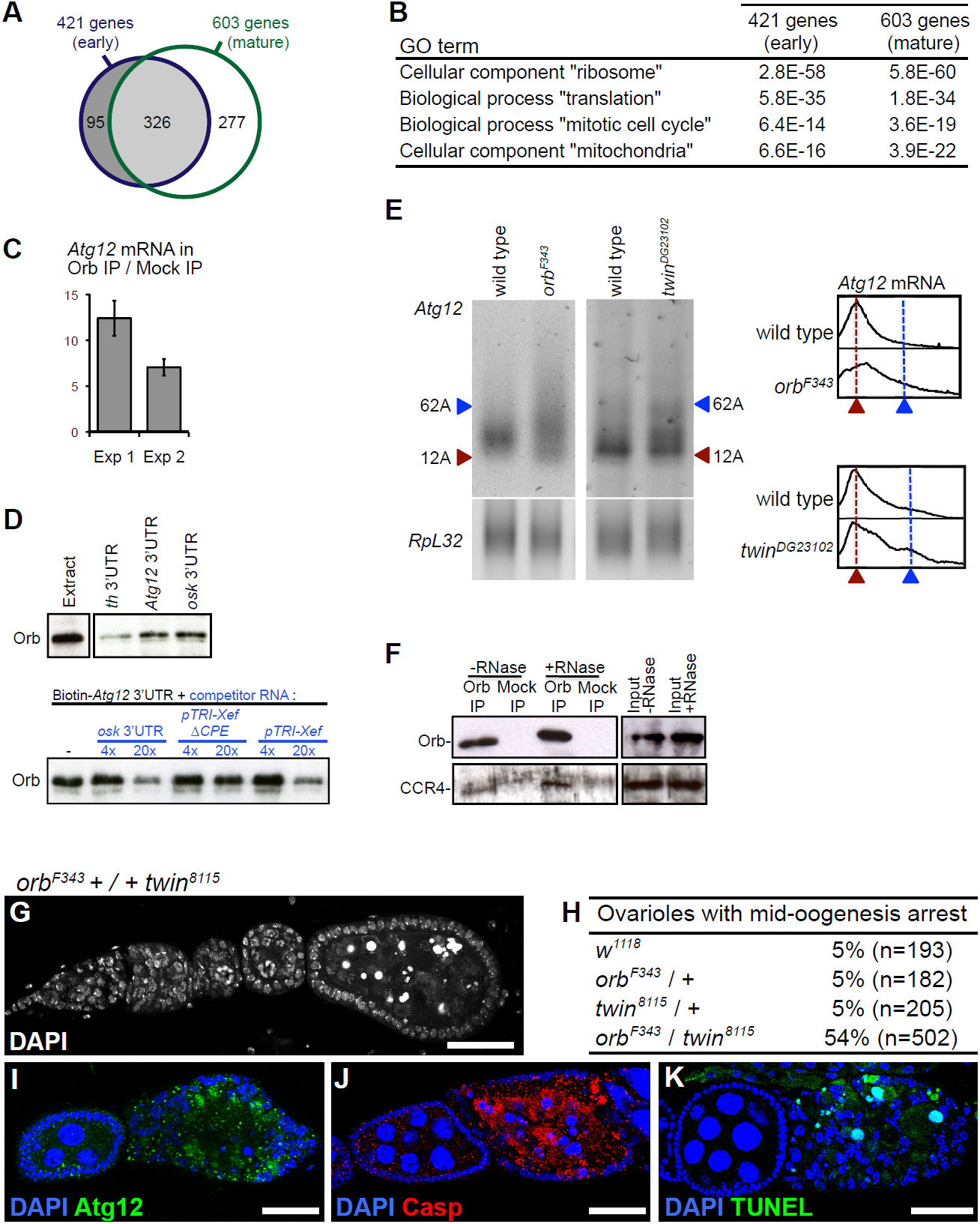
*Atg12* mRNA is a direct target of Orb. (A) Venn diagram of mRNAs significantly enriched in Orb versus mock RIP, from early (blue outline) and mature (green outline) ovaries. (B) Gene ontology term enrichment using DAVID. (C) RT-qPCR experiments showing enrichment of *Atg12* mRNA in Orb RIP versus Mock RIP. Mean ratios of three quantifications of *Atg12* mRNA levels in Orb/Mock, normalized with *RpL32* mRNA. Error bars: SEM. Two independent experiments (Exp 1, Exp 2) are shown. (D) Biotin RNA pull-down experiments showing that *Atg12* and *osk* 3’UTRs pull down Orb more efficiently than *th* 3’UTR (top panels). Biotin RNA pull-down competition assays showing efficient competition with 20X of unlabeled *osk* 3’UTR or *pTRI-Xef* which contain CPEs, and the lack of efficient competition with 20X of unlabeled RNA fragment devoid of CPE (*pTRI-XefΔCPE*) (bottom panel). (E) PAT assays showing elongated poly(A) tails of *Atg12* mRNA in *orb*^*F343*^ and *twin*^*DG23102*^ early ovaries. *RpL32* mRNA was used as a negative control. The profiles of *Atg12* PAT assays using ImageJ are shown. (F) Orb immunoprecipitation in early ovaries showing CCR4 co-precipitation, both in the absence (-RNase) and in the presence (+RNase) of RNase A and micrococcal nuclease. (G) DAPI staining of a *orb*^*F343*^ +/+ *twin*^*8115*^ ovariole showing mid-oogenesis arrest. (H) Quantification of ovarioles with mid-oogenesis arrest in control and *orb*^*F343*^ +/+ *twin*^*8115*^ females. (I-K) *orb*^*F343*^ +/+ *twin*^*8115*^ ovarioles labeled with DAPI (blue) and anti-Atg12 (green) (I); DAPI (blue) and anti-cleaved caspase 3 (Casp, red) (J); DAPI (blue) and TUNEL (green) (K). Scale bars: 20 μm.

Among the Orb mRNA early targets that might be involved in cell death, we identified *Autophagy-specific gene 12* (*Atg12*) functionally annotated for “autophagic cell death”. Since autophagy is thought to contribute to cell death in *Drosophila* oogenesis and *Atg12* encodes an effector of autophagosome formation (Gorski et al., 2003; Scott et al., 2004), we focused on this mRNA and studied its potential direct regulation by Orb.

### *Atg12* mRNA is a direct target of Orb

Independent Orb RIP and quantification of *Atg12* mRNA levels by RT-qPCR confirmed the presence of *Atg12* mRNA in complex with Orb (Figure 3C). The presence of two potential CPEs in the *Atg12* 3’UTR (Figure S3) led us to use RNA pull-down assays to address a potential direct binding of Orb to *Atg12* 3’UTR. *oskar* (*osk*) 3’UTR known to interact with Orb (Chang et al., 1999) was used as a positive control, and the 3’UTR of *thread* (*th*) which encodes DIAP1 protein was used as a negative control. Both *Atg12* and *osk* 3’UTRs were able to pull down Orb protein from an ovarian extract, while *th* 3’UTR was not to the same extent (Figure 3D). Competition assays were used to test the binding specificity of Orb to *Atg12* 3’UTR. *osk* 3’UTR and the *TRI-Xef* RNA were used as CPE-containing competitors, and *TRI-XefΔCPE* as a competitor which did not contain CPE. Unlabeled competitor RNAs were added in excess to the binding reactions (4X or 20X). The presence of 20X CPE-containing competitor RNAs substantially decreased the binding of Orb to *Atg12* 3’UTR, whereas the non-CPE competitor did not (Figure 3D). These results are consistent with the direct binding of Orb to *Atg12* 3’UTR.

### Orb and CCR4 repress *Atg12* mRNA translation through deadenylation

We addressed whether Orb was involved in the control of *Atg12* mRNA poly(A) tail lengths using PAT assays. In contrast to the shorter poly(A) tails that were reported in *orb* mutants for several mRNAs (Benoit et al., 2005; Castagnetti and Ephrussi, 2003; Juge et al., 2002), we found that *Atg12* mRNA had elongated poly(A) tails in *orb* mutant early ovaries (Figure 3E), suggesting a role of Orb in *Atg12* mRNA poly(A) tail shortening that could lead to its translational repression. The role of Orb as a translational repressor of *Atg12* mRNA was confirmed by the increased levels of Atg12 protein in *orb* mutant pseudo-egg chambers compared to wild-type (Figure 4A, B).

**Figure 4.**
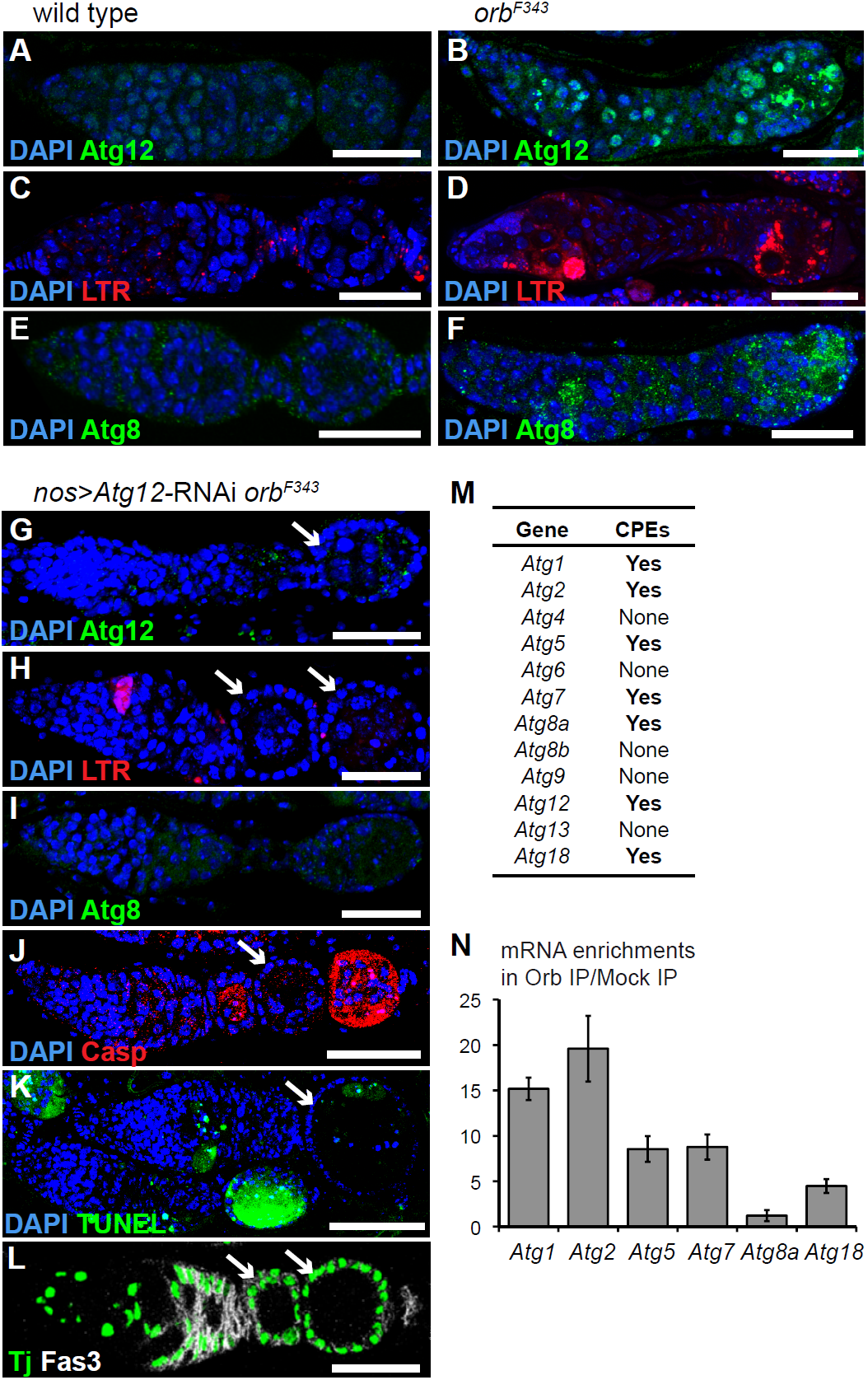
*orb* regulates autophagy. (A-L) Wild-type (A, C, E), *orb*^*F343*^ (B, D, F) and *UASp-TRIP-Atg12 orb^F343^/nos-Gal4 orb*^*F343*^ (*nos > Atg12-RNAi orb*^*F343*^) (G-L) ovarioles labeled with DAPI (blue) and anti-Atg12 (A, B, G); DAPI (blue) and Lysotracker (LTR, red) (C, D, H); DAPI (blue) and anti-Atg8 (E, F, I); DAPI (blue) and anti-cleaved caspase 3 (Casp, red) (J); DAPI (blue) and TUNEL (green) (K); anti-Tj (green) and anti-Fas3 (white) (L). White arrows indicate rescued egg chambers. Scale bars: 20 μm. (M) Presence/absence of CPEs in 3’UTRs of autophagy-specific genes. (N) RT-qPCR experiments showing enrichment of *Atg* mRNAs in Orb RIP. Mean ratios of three quantifications of *Atg* mRNA levels in Orb/Mock, normalized with *RpL32* mRNA. Error bars: SEM.

While a function of Orb in translational repression has not been reported, CPEB has been shown to act as a translational repressor prior to its action in cytoplasmic polyadenylation, and the presence of a deadenylase in the CPEB complex appears to contribute to this repressor function (Hosoda et al., 2011; Kim and Richter, 2006). We therefore analyzed the potential role of the CCR4-NOT deadenylation complex in Orb-dependent poly(A) shortening of *Atg12* mRNA. PAT assays of *Atg12* mRNA in ovaries mutant for *twin*, the gene encoding CCR4 deadenylase (Temme et al., 2004) revealed longer than wildtype poly(A) tails, indicating a role of CCR4 in shortening *Atg12* mRNA poly(A) tails (Figure 3E). Co-immunoprecipitation experiments used to determine if Orb and CCR4 could be in complex in early ovarian stages showed that Orb was able to co-precipitate CCR4 in the presence or the absence of RNAs (Figure 3F), indicating that both proteins are part of the same complex. Importantly, we also uncovered a strong genetic interaction between *orb* and *twin*. Whereas single heterozygous *orb* or *twin* mutant females displayed normal oogenesis, double *twin orb* heterozygous mutant females showed about 50% of ovarioles arrested at mid-oogenesis (Figure 3G, H). Arrested egg chambers expressed Atg12 protein, consistent with Orb and CCR4 acting together to repress *Atg12* mRNA translation (Figure 3I). Moreover these egg chambers underwent cell death as indicated by the expression of cleaved caspase 3 and staining by TUNEL assays (Figure 3J, K).

These data strongly suggest that Orb acts with CCR4 to repress *Atg12* mRNA translation by poly(A) tail shortening.

### Orb represses autophagy and cell death during oogenesis

Atg12 protein expression in *orb* mutant ovaries correlated with the induction of autophagy indicated by the presence of autophagosomes visualized using the Lysotracker marker (100% (n > 100) of *orb^F343^* ovarioles) and punctate staining of Atg8 protein (Barth et al., 2011) (Figure 4C-F). To address the functional importance of *Atg12* mRNA regulation by Orb in autophagy and cell death, we reduced *Atg12* expression in *orb* mutant ovaries using RNAi. Germline expression of *Atg12*-RNAi in *orb^F343^* mutant ovaries reduced the amounts of Atg12 protein, thus validating the RNAi transgene (Figure 4G). This led to decreased autophagy visualized with Lysotracker and Atg8-labeled autophagosomes, showing that *Atg12* expression had an important role in autophagy induction (Figure 4H, I). Strikingly, reduced expression of *Atg12* resulted in a strong rescue of egg chamber formation in *orb^F343^* mutant ovaries, with up to 62% of ovarioles able to produce egg chambers surrounded by follicle cells expressing Tj and Fas3 normally (Figures 2G, 4J-L). Staining with anti-cleaved caspase 3 and TUNEL showed that cell death was reduced in these egg chambers (62% (n = 47) and 72% (n = 18) of rescued egg chambers with no or weak anti-cleaved caspase 3 and TUNEL staining, respectively) (Figure 4J, K).

These results demonstrate that Atg12 expression plays an important role in the induction of autophagy in oogenesis and that Orb acts to directly repress *Atg12* mRNA translation thereby preventing autophagy and, to some extent, cell death.

Of the twelve autophagy-specific genes found in the *Drosophila* genome, seven (including *Atg12*) contained CPEs (UUUUAAU, UUUUAU, UUUUACU or UUUUAAGU) in their 3’UTR (Figure 4M, Figure S3). We used Orb RIP and RNA quantification by RT-qPCR to analyze the presence of these seven mRNAs in Orb RNP complexes. Six out of the seven were found enriched in Orb RIP suggesting that Orb might globally co-regulate the autophagic pathway (Figures 3C, 4N).

## Conclusion

The role played by autophagy and *Atg* genes in a wide range of biological processes is now emerging (Boya et al., 2013). Post-translational modulation of Atg proteins is recognized as an important mode of regulation of autophagy and crosstalk with other cellular processes, in response to cellular and environmental cues. Here, we add translational control as another key mechanism of regulation of autophagy. We have demonstrated the direct regulation of *Atg12* mRNA by Orb: Orb represses *Atg12* mRNA translation through its deadenylation by CCR4, thus preventing autophagy.

Autophagy is highly regulated by the levels of nutrients. During *Drosophila* oogenesis, autophagy is activated upon starvation or inhibition of the insulin/TOR signaling pathway, through the upregulation of Atg protein levels (Barth et al., 2011). On the other hand, Orb is part of the highly dynamic RNP granules defined as P (processing) bodies that are essential for translational regulations in germ cells (Weil et al., 2012). These P bodies (including Orb) undergo massive reorganization upon reduction of nutrient availability (Snee and Macdonald, 2009). Therefore, an intriguing implication from our data is that Orb, potentially with other translational regulators within P bodies, would act as a sensor of environmental cues to regulate autophagy. Consistent with this, we found that decreasing *orb* gene dosage increased cell death induced by amino-acid starvation (Figure S4). In agreement with the implication of CPEB proteins in interpreting environmental conditions, such a function has been proposed for CPEB1 in the regulation of glucose homeostasis in mouse liver (Alexandrov et al., 2012).

Translational regulation of autophagy might have a dramatic impact in many contexts, including in the expanding group of degenerative diseases involving RNA granules. Pathological RNA granules that form in neurodegenerative disorders have been proposed to be targeted by autophagy (Buchan et al., 2013; Ramaswami et al., 2013) which is conversely altered in some neurodegenerative diseases (Rubinsztein et al., 2012). In these disorders, translational deregulation of autophagy through affected RNA granules may induce a positive-feedback loop leading to enhanced production of pathological RNA aggregates.

## Experimental Procedures

### *Drosophila* stocks and genetics

Fly stocks used in this study and clonal analysis are described in Supplemental Experimental Procedures.

### Fluorescent labeling and immunostaining

Primary antibodies for immunostaining and procedures for fluorescent labeling are described in Supplemental Experimental Procedures.

### Immunoprecipitations and RNA analyses

Immunoprecipitations were performed as described (Zaessinger et al., 2006) and were followed by either RNA extraction and RT-PCR or by western blots as detailled in Supplemental Experimental Procedures. Poly(A) tail length analysis by PCR (PAT assay) and RT-qPCR using the LightCycler System (Roche Molecular Biochemical) were performed as described previously (Benoit et al., 2005; Benoit et al., 2008; Zaessinger et al., 2006) using primers listed in Supplemental Informations.

### RNA pull-down assays

UTP-biotinylated RNAs and unlabeled competitor RNAs were synthesized using T7 RNA polymerase on PCR fragments synthesized from genomic DNA with primers indicated in Supplemental Informations. RNA pull-down experiments were performed as published previously (Besse et al., 2009). For each experimental point, 20 µL of ovarian extract from 20 one day-old females were used.

## Acknowledgements

We are very grateful to A. Bergmann, L. Cooley, A. Ephrussi, J.R. Huynh, P. Lasko, R. Lehmann, K. McCall, R.S. Hawley, A. Spradling and D. StJohnston, for their gifts of fly stocks and antibodies, and to R. Mendez for the CPE identification software. We thank N. Lautredou-Audouy, Julien Cau, Julio Mateos-Langerak and Amélie Sarrazin at the MRI RIO imaging for excellent assistance with confocal microscopy. We thank B. Mollereau, J.M. Dura, and members of the Simonelig lab for advice during the course of this work. This work was supported by the CNRS UPR1142, ARC, ANR, and FRM.

